# Metabolic complementation in bacterial communities: necessary conditions and optimality

**DOI:** 10.1101/064121

**Authors:** Matteo Mori, Miguel Ponce-de-León, Juli Peretó, Francisco Montero Montero

## Abstract

Bacterial communities may display metabolic complementation, in which different members of the association partially contribute to the same biosynthetic pathway. In this way, the end product of the pathway is synthesized by the community as a whole. However, the emergence and the benefits of such complementation are poorly understood. Herein we present a simple model to analyze the metabolic interactions among bacteria, including the host in the case of endosymbiotic bacteria. The model considers two cell populations, with both cell types encoding for the same linear biosynthetic pathway. We have found that, for metabolic complementation to emerge as an optimal strategy, both product inhibition and large permeabilities are needed. In the light of these results, we then consider the patterns found in the case of tryptophan biosynthesis in the endosymbiont consortium hosted by the aphid *Cinara cedri.* Using in-silico computed physicochemical properties of metabolites of this and other biosynthetic pathways, we verified that the splitting point of the pathway corresponds to the most permeable intermediate.

## Introduction

Species that coexists on time and space form complex networks of interactions which are shaped by abiotic and biotic factors (Faust and Raes, 2012; Seth and Taga, 2014). The association between different species can be analyzed from the perspective of the metabolic interactions. In this context, the possible nutritional interactions within different organisms can be grouped as follows: 1) competition for limiting nutrients; 2) syntrophy, *i.e.* the cooperation emerging as each of the partners gain by the metabolic reactions of the other, for example when one organism consumes a product of some other organism, allowing the producer to drive an originally energetically unfavorable metabolic process; 3) commensalism or nutrient cross-feeding, in which the presence of an organism that over-produces an essential nutrient enables auxotroph organisms to survive. While in the case of competition one of the organism will inevitably exclude the other (Hardin, 1960), in the cases of syntrophy and cross-feeding, the interaction will tend to stabilize the coexistence of both species, otherwise competitive. In particular, when the exchange of nutrients or precursors is bidirectional and thus, is beneficial for both partners, the nutritional interdependence will lead, in most of the cases, to a co-evolutionaryprocess, the most striking example being the emergence of symbiotic associations (Douglas, 1998; Germerodt et al., 2016; Hansen et al., 2007).

In the context of the evolution of endosymbiotic bacteria the concept of functional complementation is widely used to describe the complex patterns of nutritional interactions between the members of the association. This concept has been applied at least with two different meanings. On one hand, it can be used in the sense of nutritional cross-feeding between the endosymbiont bacteria and its host, *i.e.* there are exchanges between the symbionts and/or the hosts of some essential compounds (*e.g.* amino acids, vitamins, etc.) (López-Sánchez et al., 2009]; Macdonald et al., 2012; Russell et al., 2014; Wu et al., 2006). On the other hand, the concept is also used to refer to a more complex scenario, in which the patterns of gene loss and retention in each symbiont have lead to an inter-pathway genomic complementation (Van Leuven et al., 2014). This means that during the co-evolutionary process, a subset of genes coding a certain metabolic pathway are retained in one symbiont, while the other genes are retained in the other(s) member(s) of the association. In other words, the coding genes of one or more pathways end up distributed among different genomes. In the present paper we will use the second definition.

Events of metabolic complementation, as defined above, are commonly found between endosymbiont bacteria and their hosts, where some metabolic functions are distributed between them (Baumann et al., 2006; Rao et al., 2015; Zientz et al., 2004). For example, in different obligate endosymbiont hosted by sap feeding insects, transaminase activities, a critical step in the biosynthesis of amino acids, have been found to be lost (Baumann et al., 2006; Jiang et al., 2012; McCutcheon and von Dohlen, 2011; Rao et al., 2015; Zientz et al., 2004). In some of these cases, the transaminase activities have been found to be encoded in the host genome, indicating that the synthesis of amino acids requires the combined metabolic capabilities of both members of the association (Hansen and Moran, 2011; Russell et al., 2013). Moreover, it has been suggested that the host can regulate the production of amino acids by controlling the supply of key precursors, such as glutamine (Poliakov et al., 2011; Price et al., 2014; Russell et al., 2014). Other examples of metabolic complementation include the menaquinol biosynthetic pathway in the cockroach endosymbiont *Blattabacterium cuenoti,* where mevalonate, one of the precursors, is provided by the insect host, while the symbiont provides vitamins and other essential compounds (Ponce-de-Leon et al., 2013).

Furthermore, events of metabolic complementation have also been found between the members of endosymbiont consortia in different insects, *i.e.* certain metabolic capabilities are distributed among the different endosymbiotic bacteria living in the same host. A particularly interesting example is the case of the endosymbiont consortium of the cedar aphid *Cinara cedri*. In this system, two different bacteria, *Buchnera aphidicola* BCc and *Ca.* Serratia symbiotica SCc (hereafter referred to as *S. symbiotica* SCc), coexist (Burke and Moran, 2011; Lamelas et al., 2011; Pérez-Brocal et al., 2006). In this system, most of the biosynthetic processes, *e.g.* biosynthesis of some amino acids, are being performed by only one member of the consortium, and thus each member has to rely on the other for specific nutrients. Other (non-essential) amino acids are, instead, only provided by the host, which also provides the critical transamination steps in the synthesis pathways of the symbionts. Tryptophan biosynthesis is a notable exception. This biosynthetic pathway is split in two parts, each operating in one member of the consortium. Therefore, in this case, the biosynthesis of tryptophan requires the presence of both endosymbionts. Surprisingly, a case of convergent evolution has been also found in the symbiotic system of the psyllid *Heteropsylla cubana* (Martíez-Cano et al., 2015). In this second case, the primary symbiont *Ca.* Carsonella ruddii encodes the first step of the pathway, whereas the secondary symbiont, which has lost almost all the genes for the biosynthesis of essential amino acids, still encodes the remaining genes for complementing the biosynthetic pathway (Sloan and Moran, 2012). More complex scenarios have been described in a cicada species of the genus *Tettigades* harboring endosymbionts. In this system it was found that the endosymbiont *Ca.* Hodgkinia cicadicola has evolved into two cytologically distinct species, partitioned into discrete cells, but which are metabolically interdependent (Van Leuven et al., 2014).

Most of the described complementation events have been identified through genomic analyses. However, these studies do not address the possible advantages or disadvantages of the observed metabolic design. Particularly, metabolic complementation presents certain biophysical problems regarding the splitting a metabolic pathway into different organisms. For example, there is the question of how the intermediate metabolites are exchanged between the endosymbiont and its host, or between the different members of a consortium, (*e.g. B. aphidicola* BCc and *S. symbiotica* SCc in the cedar aphid). This question becomes even more puzzling when considering that obligate endosymbionts have a very small repertory of genes coding for transporters (Charles et al., 2011). In addition, intermediate metabolites in biosynthetic pathway do not usually have associated transporters, which suggest diffusion as the most plausible mechanism for the exchanges with the surrounding environment.

Another question is how the endosymbionts adapt their pathways to satisfy the needs of the host and of the other symbiotic endosymbionts. As the symbiotic relationships are established, bacteria overproduce nutrients needed by the host. The flux through the corresponding biosynthetic pathway can be increased by acting on several properties of the enzymes (Kacser and Burns, 1973). The catalytic constant of the enzymes, and their affinities for the substrates, can be selected in order to yield larger fluxes. For instance, it has been shown that a selective pressure of improving the enzyme's efficiencies can affect lateral gene transfer in endosymbionts (Ringemann et al., 2006). A more straightforward way to increase reaction fluxes it to increase the enzyme levels. Although, metabolic processes are regulated at different levels (*e.g.* transcription), a common feature of obligate endosymbionts is the aparent absence of transcriptional regulatory mechanisms (Moran and Bennett, 2014; Wilcox et al., 2003). Therefore, enzyme levels can be increased either tuning the translation/transcription efficiency of the gene, or by modifying the gene copy number, inserting additional copies of the gene in the chromosome or using plasmids. For instance, many *Buchnera* strains, producing tryptophan for their hosts, possess multiple copies of the enzyme anthranilate synthase, which is considered to be a limiting step of the tryptophan biosynthetic pathway (Lai et al., 1994). The activity of the enzymes is also regulated by the presence of inhibitors or activators. In most biosynthetic pathways, flux is negatively regulated (through allosteric inhibition) by the final product of the pathway on the first reaction. One might also ask what is the role of feedback inhibition in systems like the *C. cedri* consortium, in which the metabolite inhibiting the pathway is not produced in the same cell. Does the division of labor operated by the members of the consortium effectively reduce the amount of allosteric inhibition and increase the production rate of tryptophan?

The very presence of metabolic complementation between symbionts and their hosts, and especially among the members of an endosymbiont consortium,rises many evolutionary questions: does metabolic complementation provides any selective advantage for the whole system? And, if it does, which is the evolutionary advantage provided by such strategy? Studies have been performed in order to elucidate if organisms exhibiting cross-feeding interactions do possess any selective advantage with respect to free-living organisms (Germerodt et al., 2016; Pfeiffer and Bonhoeffer, 2004). Microbial coexistence has been also explored by using thermodynamic and kinetic models (Großkopf and Soyer, 2016). At any rate, understanding the evolutionary and mechanistic properties of natural microbial communities might allow the design of synthetic ones with potential industrial interest (Großkopf and Soyer, 2014).

## The model

### A kinetic model of two parallel unbranched pathways with negative feedback inhibition

We developed a kinetic model to study metabolic complementation endosymbiont consortium of two different bacteria, when they are allowed to exchange intermediate metabolites of a hypothetical biosynthetic pathway. The model describes a consortium consisting of two different endosymbiotic bacterial species and their host (Fig 1). Each of the two bacterial populations is represented by a large number ( *N_i_* with *i* = 1,2) of identical cells, so that in practice it is sufficient to consider two different compartments, modeling a single cell of each bacterial strain, enclosed in a larger compartment, which allows for exchanges of metabolites between cells and the host. Both endosymbionts code for the same hypothetical biosynthetic pathway, which allows the conversion of a precursor *S* (produced by the bacteria), to an end product *P*, which is essential for any of the three members of the consortium. The biosynthetic pathway is a linear chain composed by two reactions (see Fig. 1B), with the precursor *S* being first converted into an intermediate metabolite *X*, and then *X* being converted to the final product *P*.

**Figure 1.**
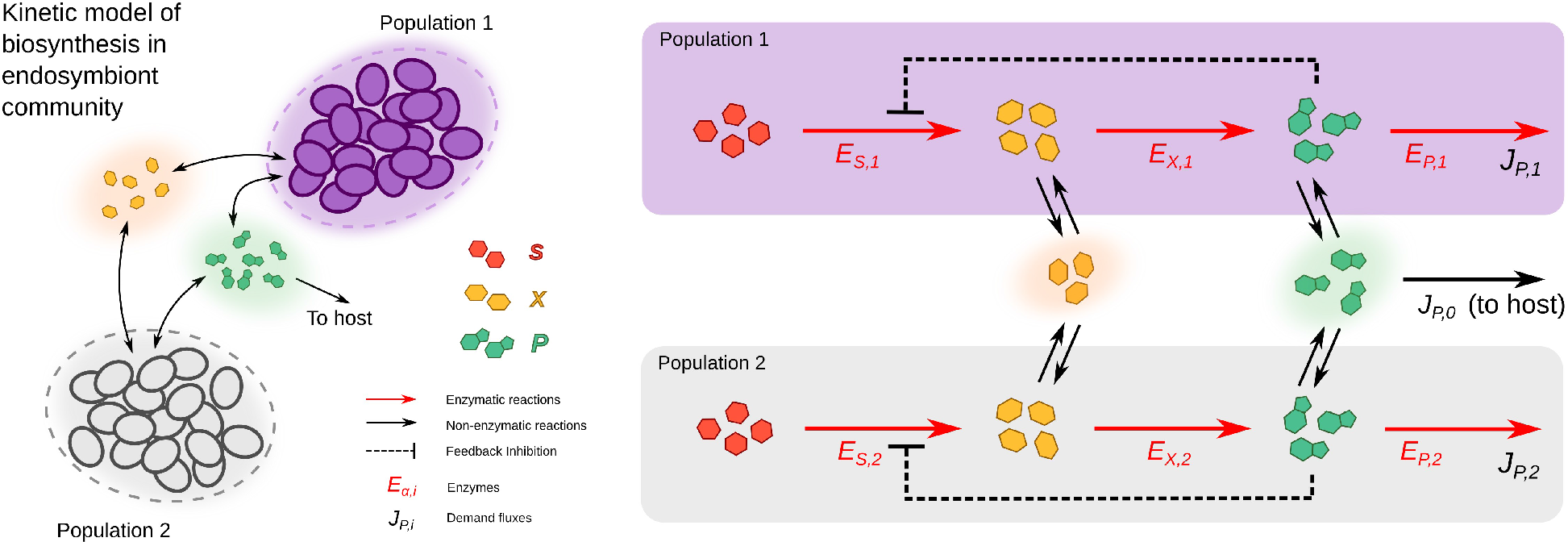
*Kinetic model of metabolic complementation.* *Two cell types (purple and gray) can both potentially produce the product *P* starting from a substrate *S*, passing through an intermediate metabolite *X*. The enzymes can be freely allocated to the pathway’s reaction and consumption of *P*. The intermediate ( *X*) and final ( *P*) metabolites can cross the cell membranes by diffusion. Both bacteria and the host need the final product to grow; leading to consumption rates *J_P,0_*, *J_P,1_* and *J_P,2_*. In the case of the Buchnera/Serratia/Cinara consortium, the pathway is completely split across the two populations ( *E_S,2_*and *E_X,1_*. are absent).*

The intermediate metabolite *X* and the final product *P* can freely diffuse outside the cells (Fig. 1B). Moreover, since the product *P* is considered essential for both endosymbionts and its host, three demand fluxes are also introduced to represent the incorporation of *P* to the respective biomasses of the three member of the consortium. Finally, product inhibition has been suggest to be present in some endosymbionts, *e.g.* anthranilate synthase of *B. aphidicola* shows a highly conserved allosteric binding site (Lai et al., 1994), albeit in other enzymes like prephenate dehydratase the inhibition could be mitigated by point mutations in the allosteric site (Jiménez et al., 2000). Thus, the model includes a negative feedback inhibition mechanism that regulates the production of *P*.

The two reactions in the biosynthetic pathway, *S → X* and *X → P*, are modeled using reversible Michaelis-Menten kinetics (Noor et al., 2013); we name the corresponding fluxes *V_S_* and *V_X_*, respectively. Moreover, in order to include the effect of product inhibition on the first step ( *S → X*), a term representing such effect was introduced in the kinetic equation of the first reaction (Cornish-Bowden, 2013). There are diverse mechanisms of enzyme inhibition, including allosteric interactions, but in this approach we have analysed one of the simplest situations and the kinetics of the inhibition was considered, in a first place, to be uncompetitive, although the case of competitive inhibition is discussed later. Thus, the kinetic law of the first reaction *S →X*, with flux *V_S_*, is the following: 
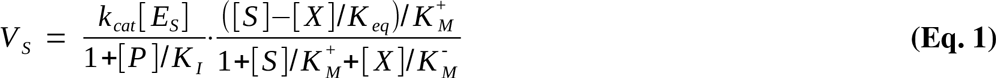

where *k_cat_* is the turnover number, or catalytic constant, of the reaction, 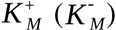 is the forward (backward) Michaelis constant, *K_eq_* is the equilibrium constant, and *K_I_* is the inhibition constant. It is worth to note that in the limit 1**/***K*_I_ →0 0 the reversible Michaelis-Menten kinetic law, without inhibition, is recovered. The kinetic for the second reaction ( *X → P*, with flux *V_X_*) is analogous to (Eq. 1) but without the inhibition term ([*P*]**/** *K_i_*).

In order to allow the flux of metabolites between the members of the consortium, the metabolites *X* and *P* are allowed to be exchanged with the surrounding environment. Endosymbiotic bacteria usually lack transport systems (Charles et al., 2011); thus, the exchanges of *X* and *P* are modeled using first-order, non-saturating, reversible kinetic laws. The fluxes *U*_α_ take the following form:

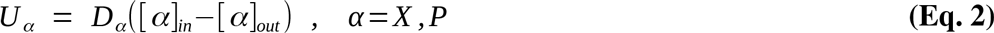

where *D*_α_ are related to membrane permeability and diffusion constants of the metabolites (Laidler and Shuler, 1949). Note that the fluxes are assumed to be positive when the metabolites are excreted by the cells. On the other hand, if there were any kind of protein-mediated transport, the corresponding membrane proteins should also have been taken into account. Finally, the product *P*is assumed to be consumed during growth by the three members of the consortium, with consumption fluxes in each compartment. For the bacterial cells, we model the consumption of the metabolite *P* using an irreversible Michaelis-Menten reaction: 
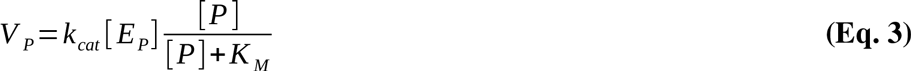

The fluxes *V_P,i_* (the index *i*= 1,2 corresponding to the cell type) are constrained to be larger than some fixed flux *J_P,i_*. The extracellular product is instead consumed by the host. It is useful to normalize this rate to the size of the bacterial populations *N=N*_1_**+***N*_2_, so that the consumption rate equals *d[P]*_out_**/**dt= – *NJ*_P,0_,. The three parameters *J_P,0_* and *J_P,i_* thus set the demand of the metabolite *P* from the bacteria or the host.

Finally, the size of the populations of the two bacteria naturally influence both the total protein concentration and the magnitude of the transport fluxes between the cells and the host, and cannot be disregarded. It is convenient to introduce the normalized populations n_1_=*N*_1_**/** *N* and *n_2_= N_2_**/** N*, with *N_1_**+**N_2_= N*. We will therefore first show results obtained by fixing the cell relative populations to be n_1_ = n_2_ =1**/**2, and then exploring the case in which the two population fractions differ. When the whole model is considered, the balance equations for all the metabolites of the system can be written as follows:

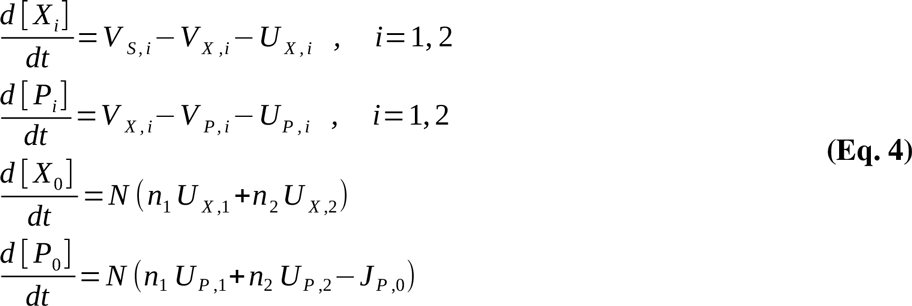

where *N* corresponds to the total number of cells in both populations. Is worth to note that the equations for intracellular metabolites are symmetric for the two species, while the balance equations of the extracellular metabolites depend on the size of each symbiont population and, in the case of *P*_0_, on the host demand flux *J_P,0_*. The substrate *S* is a precursor produced by each of the two cell types, and its concentration is assumed to be fixed (Heinrich and Schuster, 1996). As a consequence, the balance equation of *S* is not included in the model. For the sake of simplicity, in the following we will set all kinetic parameters to one, unless stated otherwise. Also, we will set the population fractions to *n*_1_=*n*_2_=1/2 and the concentration of the substrate to [*S*] = 10

### The optimization criteria

Endosymbiont bacteria involved in nutritional interactions overproduce some essential products, such as essential amino acids or vitamins, and share them with their host (and in some cases with other endosymbionts) (Douglas, 1992). On the other hand, as discussed in the introduction, such over-production can be ensured by tuning the enzyme concentrations along the pathway, even if end-product inhibition may not allow to tune the flux easily (Baumann et al., 1999; Lai et al., 1994; Rouhbakhsh et al., 1997). Accordingly, improvements in the efficiency of over-productions, in terms of the economy on protein synthesis, should provide of some advantage to the whole system (Dekel and Alon, 2005; Flamholz et al., 2013; Goelzer and Fromion, 2011; Kafri et al., 2015; Mori et al., 2016; Ponce de León et al., 2008; Shachrai et al., 2010). Thus, in this work we will focus on the trade-off between the maximization of the flux through a pathway and the efficient allocation of the proteins. Using constraint-based optimization this trade-off can be modelled as two complementary optimization problems: (i) maximization of the flux subject to a constraint on the maximum amount of total enzyme concentration; and (ii) minimization of enzyme concentrations subject to a constraint (lower bound) on the flux of interest, which ensure that a certain demand is satisfied.

Here in, we will adopt the second formulation, *i.e.* enzyme minimization subject to a certain demand flux of the product *P*, although one can show that both formulations yield completely analogous results (see Supplementary Text S1). The optimization problem is defined in the following way. Given the demand fluxes *J_P,i_*, *i* = 1,2:

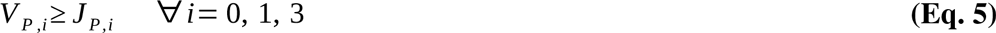

we compute the metabolite and enzyme concentrations which minimize the total enzyme concentration (normalized by the total bacterial population), defined as follows:

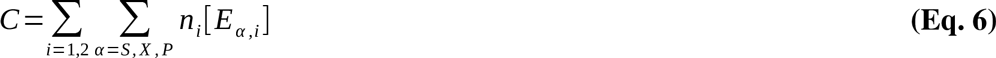

where the cost or objective function *C* correspond to the total amount of protein invested by both cell types in order to produce an amount of *P* enough to satisfy the demand required by the three members of the consortium. All the intermediate metabolites are subject to the steady-state condition, which is imposed by setting to zero the time derivatives in (Eq. 4). Accordingly, each solution will consist of a set of metabolites and enzymes concentrations, from which one can compute the optimal fluxes using the kinetic laws. A complementary formulation of the optimization problem is possible, *i.e.* maximization of the flux subject to a constraint on the maximum amount of total enzyme concentration. This other approach can be shown to be completely equivalent to the “enzyme” minimization approach, as shown in Supplementary Note 1.

In order to highlight the patterns of proteome allocation between the cell populations, we introduce the measure of enzyme asymmetries *A*_α_, defined for each enzyme α as follows:

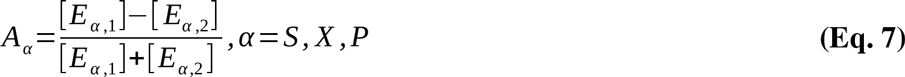

This quantity reflects the relative difference in the concentration of each enzyme between the two different endosymbiont. Thus, *A*_α_ will be zero if the enzyme concentrations in the two cells are the same, while it will equal either 1 or-1 if one cell type completely lacks the corresponding enzyme. In particular, if the two type of cells have the same sets of kinetic parameters, one would expect that the protein asymmetries would be equal to 0, unless some metabolic interaction emerges.

## Results

### Emergence of metabolic complementation

Using the model and the optimization criteria introduced in previous sections, we now turn on the analysis of the optimal proteome allocation obtained, for different scenarios. As a starting point, we will set the host demand flux to zero, *J*_P,0_=0. As explained in Supplementary Note S1, since the total enzyme concentrations is proportional to the demand fluxes, the sum of the demand fluxes *J*_P,1_**+** *J*_P,2_is not relevant for the optimal strategy of proteome allocation, and only their ratio *J*_P,1_**/***J*_P,2_ is (as long as *J*_P,0_=0). We will therefore choose *J*_P,1_=1 for both the two cell types. (The results for different values of *J*_P,0_=0 and *J*_P,1_**/***J*_P,2_ are shown below.)

We show in Fig. 2 the optimal protein asymmetries obtained by varying the value of the diffusion constants, *D_X_* and *D_P_*, for different values of the inhibition constant *K*_I_. Given the symmetry of the problem, i.e. the reactions in the two cell types have identical kinetic parameters, we will only consider the solutions with *A*_S_≥0. In the extreme case where *D_X_*=*D_P_*=0, the two cells cannot interact in any way and no complementation can arise. However, when the metabolites are allowed to permeate, the two cell populations are in principle allowed to interact, and an asymmetric solution corresponding to metabolic complementation may emerge. If the effect of the product inhibition is too low (*e.g.* 1/*K*_I_≤0.1), then the optimal solution is always characterized by a symmetric enzyme allocation, i.e. *A_α_*=0, for all enzymes involved (Fig 2A), and for all values of the permeability constants *D_X_* and *D_P_*. Conversely, when the inhibition gets larger and larger (*e.g.*1**/***K_I_* ≥1), asymmetric solutions starts to emerge as the optimal strategy, depending on the values of the diffusion constants *D_X_* and *D_P_* (Fig 2B-D). For a given value of 1**/***K_I_*, the diffusion constants define a transition line, which divides the plane into two zones, corresponding to the range of parameter for which the symmetric or asymmetric solution is optimal.

**Figure 2.**
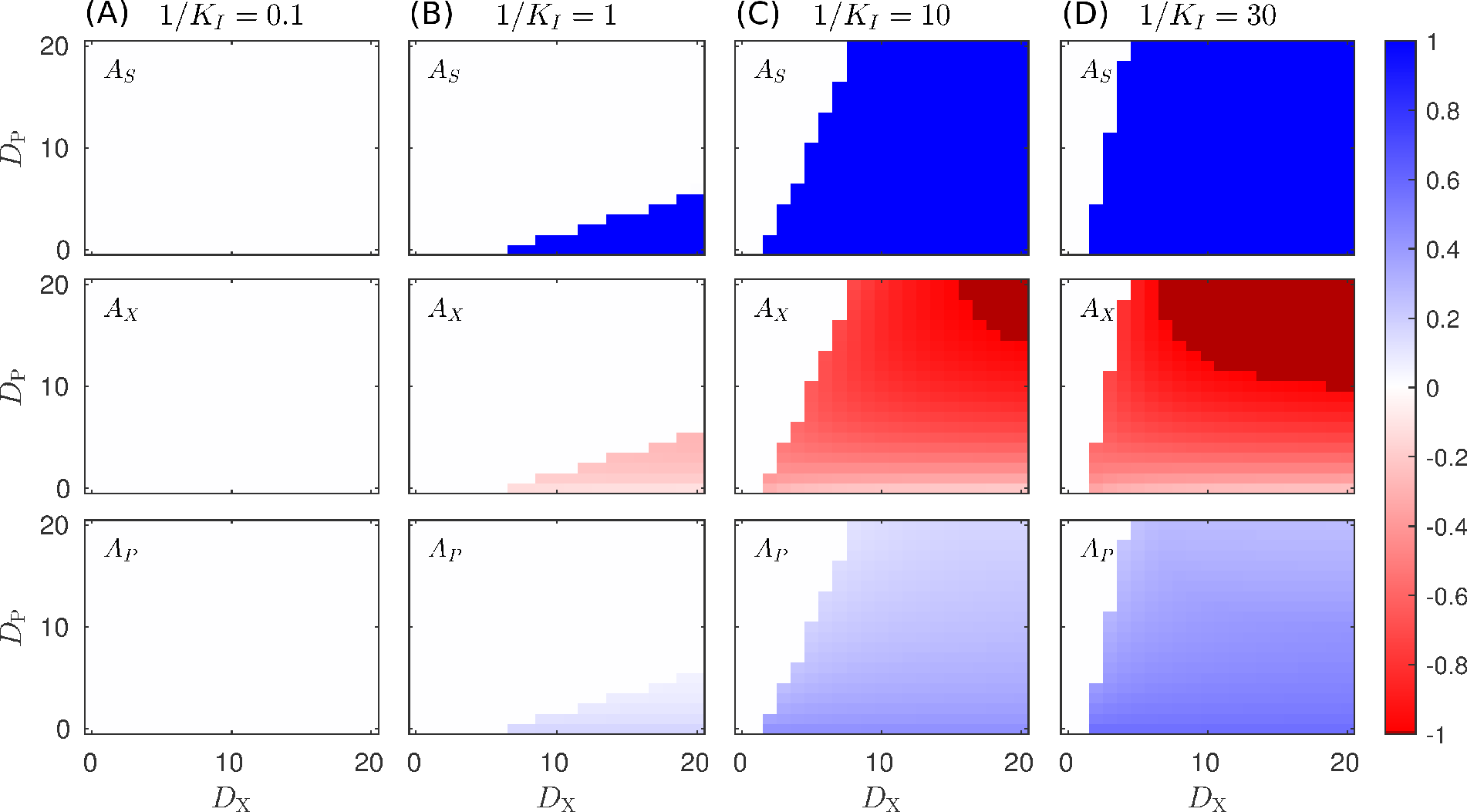
*Enzyme asymmetries A_α_ for optimal proteome allocation solutions obtained in different scenarios.* *The figure shows the optimal solution in terms of the enzyme asymmetries as a function the diffusion constants D_X_ and D_P_, and for different values of the inhibition constant, in the case of uncompetitive inhibition of P on the enzyme E_S_ (see Eq. 1), Subplots on each column, labeled A-D correspond to the solution obtained for a different values of 1**/**K_I_, whereas rows represents the different enzymes of the pathway. The darker color in the 1**/**K_I_=10 and 1**/**K_I_=30 subplot columns highlight the region in which [E_S,2_]=[E_X,1_]=0, and the pathway is completely split between the two cell types. The color scale corresponds to the asymmetry value of the different enzymes.*

When the condition favors the asymmetric strategy, in general all the enzymes exhibit certain degree of asymmetry. In particular in the case of the enzyme *E_S_*, the asymmetry only takes the extreme values, *A_S_* = 0 or |*A_S_*| = 1, whereas for the other two enzyme the degree of asymmetry depends on the diffusion constants. Interestingly, we find that large values of *D_X_* promote the asymmetry, while for large values of *D_P_* the symmetry is restored. In other words, asymmetric solutions arise when *D_X_ **/** D_P_* is large enough. Summarizing, asymmetric proteome allocation becomes an optimal strategy in the presence of byproduct inhibition and when the product diffuses slower than the intermediate metabolite *X*. If *P* diffuses too fast, the concentrations within both cell types will be roughly the same, and therefore the inhibition cannot be by-passed. Conversely, if *P* diffuses much slower than *X*, its concentration on the cell coding for the first enzyme (i.e. *E_S_*) will be low and its contribution as an inhibitor will be negligible. As shown in Fig 2, in the regions where asymmetry emerges as the optimal solution, we have the following relations for enzyme asymmetries: *A_S_* =1, *A_X_*< 0 and *A_P_*> 0. Thus, in order to have a deeper insight on the relation between diffusion and the sudden change from symmetric to asymmetric solution, we focus now on single value of the inhibition constant (1**/***K*_1_=10). We can individually study the symmetric and asymmetric solutions by imposing that *A_S_*=0 (in the symmetric case) or *A_S_*=1 (in the asymmetric case). Then, for these two cases, the optimal proteome allocation was computed for different values of the diffusion constants, *D_X_* and *D_P_*.

Fig 3A shows that the protein cost of the asymmetric strategy grows when the metabolites cannot easily diffuse across the membranes, i.e. when the values of *D_X_* and *D_P_* decrease (yellow surface/lines). On the other hand, the asymmetric strategy becomes optimal for large values of the diffusion rates. For large values of the diffusion constants we see that the asymmetric solution is optimal, with the *A_X_* enzyme asymmetry approaching the value-1 (Fig. 3B). In this conditions, the pathway is completely splitted between the two cells, with one enzyme ( *E_S_*) only present in the first cell (Fig. 3B), and the second enzyme ( *E_X_*) only present in the other cell (Fig. 3C). We see in Fig. 3D that the asymmetric condition, *A_S_* = 1, is always accompanied by an excretion of the *X* metabolite from the first cell, which is gathered by the second cell *U_X,1_*= – *U_X,2_*> 0. Interestingly, the exchange of *X* (Fig. 3D) still takes place even in the case in which *P* cannot cross the cell membrane, i.e. *D_P_*=0 (Fig. 3E). In this condition, the second population becomes a cheater, and takes advantage of the leakage of *X* from the first population. Nevertheless, if the diffusion constant *D_P_* is increased, the second cell starts sharing the product *P* with the first cell, thus establishing a metabolic complementation relationship. Finally, if *D_P_* is too large, the concentrations of the *P* metabolites are very similar in both cells, so that there is no way to reduce the effect of inhibition by partitioning the metabolic route. Therefore, the cell switches from metabolic complementation to an “neutralist” strategy, with completely symmetric enzyme allocation and no exchange of metabolites between the two cells types.

**Figure 3.**
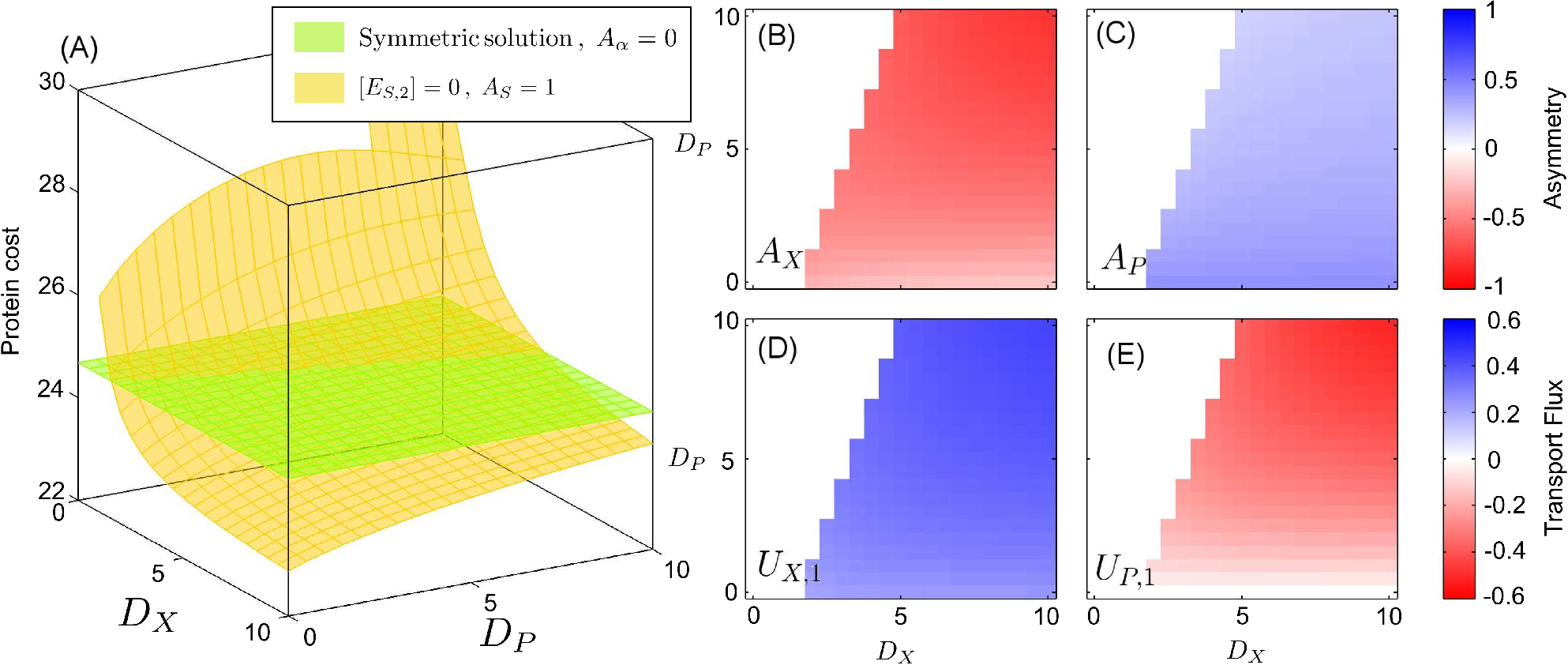
*Optimal solutions (minimum proteins) in the case of uncompetitive inhibition.* *The figure shows protein costs plotted as a function of *D_X_* and *D_P_* for a fixed value of the inhibition 1**/***K_I_*=10.(A) The cost function (that is, the total protein cost) is shown in the z-axis; the green surface, corresponding to the symmetric solution, is obtained by imposing that the two cells have identical enzyme allocation, while the yellow one is obtained imposing that one of the two cells lacks the *E_S_* enzyme. (B) proteome asymmetry of the *E_X_* enzyme; (C) Proteome asymmetry for the *E_P_* enzyme; (D) Optimal transport flux *U_X,1_*, of the *X* metabolite from cell 1 to cell 2; (E) Optimal transport flux *U_P,1_*; of the P metabolite from cell 1 to cell 2 (the negative value of the flux means that *P* is actually shuttled from cell 2 to cell 1). The color scale of subplots B-C and D-E, corresponds to the asymmetry value of the different enzymes and the sense of the transportation fluxes, respectively.*

## Assessing the impact of alternative inhibition mechanisms and different product demands

In this section we extend our results to more general settings. First, we check that our results do not depend qualitatively on the inhibition kinetics. Instead of a non-competitive mechanism (Eq. 1), we use a competitive mechanism (Cornish-Bowden, 2013) so that the concentration of the product metabolites affects the saturation of the enzyme, and not the turnover rate. Although this mechanism is more complicated than the uncompetitive case, due to the interplay between the different concentrations of the *S*, *X* and *P* metabolites, the results obtained are analogous to the simpler case of uncompetitive inhibition (see Supplementary Fig. S1). This suggests that while product inhibition is a necessary condition for the emergence of metabolic complementation, it does not depend on specific inhibition mechanisms, and the metabolic complementation strategy is favored by strong feedback inhibition.

Then, we probed the effect of different demand fluxes, both to the host (*J_P,0_*) and to the bacterial species (*J_P,1_* and *J_P,2_*,). As one increases the host demand flux *J_P,0_*, the qualitative pattern of the different optimal strategies is only slightly affected by the presence of a flux of products to the host (see Supplementary Fig. S2), even though the total amount of protein to be allocated clearly increases with the demand flux. Similarly, we consider the case where the two bacterial populations have different demands for the product *P*, for instance because of a different biomass composition. We see that an asymmetry in the demands for *P* can promote, even more, the asymmetry in the proteome allocation (see Supplementary Fig. S3); for large enough 1/*K_I_* and demand flux asymmetry ρ_J_ =(*J_P,1_*−*J_P,2_*)**/**(*J_P,1_***+***J_P,2_*), the optimal solution is characterized by a complete splitting of the pathway within the two cells, [*E_S,1_*] =[*E_X,2_*] = 0.

A similar effect is found when the two bacterial species have different populations, that is, *n*_1_≠ *n*_2_. As n_1_ is moved away from ½ (with *n*_2_=1− *n*_1_), the protein asymmetry is generally enhanced, with larger asymmetries *A_X_* (see Supplementary Fig. S4). All these results show that the complementation patterns described by the model are robust against variation in the demand fluxes and the relative population sizes.

## Sensitivity analysis

The number of kinetic parameters in the model is quite large, and it is very difficult to probe the effect of perturbations for each different combination of parameters. Therefore, we performed a sensitivity analysis, on the model with uncompetitive inhibition, by randomly perturbing the parameters, and studying how the perturbation of each parameter correlates with the insurgence of the asymmetric optimal solution (see Materials and Methods for details). We see that the parameters which promote the enzyme asymmetry most are, as expected, the inverse inhibition constant 1**/***K_j_* (reflecting the amount of inhibition), the concentration of the *S* metabolite, and the diffusion constant *D_X_* of the *X* metabolite (Supplementary Table S1). We also find that the equilibrium constant of the *S→X* reaction also positively affects the asymmetry, albeit slightly; this is because increased concentrations of the *X* metabolite allows for larger shuttling fluxes across the two cells. On the other hand, the diffusion constant *D_P_* of the *P* metabolite is found to promote the symmetry between the two cells, decreasing the average value of *A_S_*.

## Extended model with multiple intermediates

In the previous section we gave an in depth characterization of a modelin which the biosynthetic pathway is composed by two reactions. However, real biosynthetic pathways are usually composed by a larger number of reactions. In order to study how the permeability affects the emergence of a complementation, an extended version of the model was developed, which keeps the fundamental features of the its simplest version, while adding more intermediates in the pathway. The extended version of the model was obtained by introducing five intermediate metabolite, *X_1_*, …, *X*_5_, along with five enzymes *EX*_1_, …, *EX*_5_. Each enzyme *EX_i_* with the index *i* ranging between 1 and 5, catalyzes the reversible conversion of *X_i_* into *X*_i+1_ or, in the case of *EX*_5_, the product metabolite *P*. This extended model allows us to study how a long biosynthetic pathway is differently partitioned within the two cells for different permeabilities of the intermediate metabolites.

**Figure 4.**
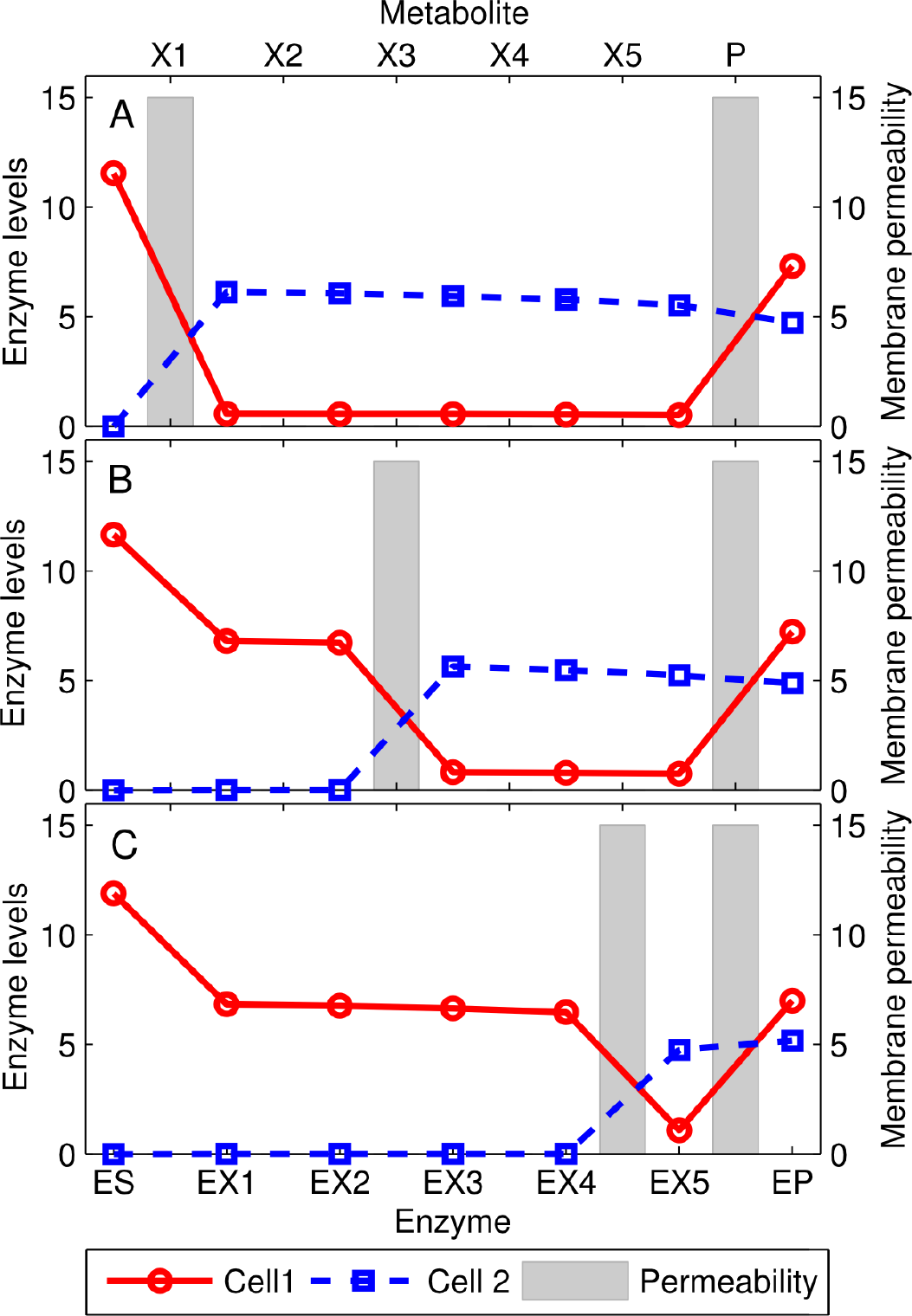
*Solutions or the extended model with 5 intermediate metabolites *X*_1_,… *X*_5_.* *The points represent the optimal enzyme levels of the two cells, while the gray bars are the membrane permeabilities ( D_i_) for the intermediate metabolites. In each panel, the P metabolite and one of the intermediate metabolites X_i_(*i*= 1, 3 and 5 in panels A, B and C, respectively) is allowed to permeate through the membrane ( *D* = 15), while all others were not ( *D*=10^-3^). In these conditions, the pathway is split in correspondence of the permeable intermediate metabolite. Other settings: *K*_eq_=4**/**3 for all reversible reactions (all but the one catalyzed by the E_P_ enzyme). Inhibition constant: *K_I_*=1**/**10; the concentration of [*S*]=20.*

In the extreme case in which only one intermediate metabolite can permeate through the cell membranes, we find that the general behavior of the extended model is totally consistent with the results obtained using the simplified version (see “Emergence of metabolic complementation” section). As shown in Fig. 4, the pathway is split in correspondence of the most permeable intermediate metabolite. The first cell converts the substrate *S* into the intermediate metabolite with large permeability, which in turn is (mostly) shuttled to the other cell and converted into the product *P*, which is then shared between the two bacteria. A wide range of behaviors are obtained when different, more complex, scenarios are considered (see Supplementary Fig. S5). For example, when all the intermediate metabolites can permeate across the membranes, the optimal solution still exhibits metabolite complementation; However, enzymes levels change smoothly along the pathway with many metabolite being exchanged, and therefore the qualitative pattern lacks a unique breaking point.

## Predicting Biological cases: The case of the tryptophan biosynthesis

In this section we discuss the well documented case of the complementation of the tryptophan biosynthetic pathway between the members of an endosymbiotic consortium in relation with the results here obtained (Gosalbes et al., 2008; Sloan and Moran, 2012). This event of metabolic complementation has been reported to be convergent in two different insects systems and evolutionary unrelated. On one hand, the complementation of the tryptophan biosynthesis has been described in the endosymbiotic consortium composed by the P-endosymbiont *B. aphidicola* BCc and the co-primary endosymbiont, *S. symbiotica* SCc, hosted by the cedar aphid (Gosalbes et al., 2008). On the other hand, the complementation has also been described in the system of the psyllid *Heteropsylla cubana,* where the coding genes for the tryptophan pathway are distributed among the genomes of two endosymbionts, *Ca.* Carsonella ruddii and a tentatively identified secondary symbiont (Sloan and Moran, 2012), thus, exhibiting the same pattern than the one described for the case of the aphid *C. cedri* (Martíez-Cano et al., 2015). It is worth to note that although aphids and psyllids belongs to the same suborder Sternorrhyncha, their corresponding endosymbionts belong to different bacteria lineages, indicating the independence of the symbiotic events (Baumann et al., 2006). This convergence suggests the existence of some evolutionary advantage of the complementation strategy. In the following we will focus on the case of *C. cedri*, because this system has been extensively characterized (Burke and Moran, 2011; Gosalbes et al., 2008; Lamelas et al., 2011; Pérez-Brocal et al., 2006).

The tryptophan biosynthetic pathway involves five enzymes, encoded by seven genes (*trpA-F*), which allows the conversion of chorismate to tryptophan (Crawford, 1989) (see Fig. 5). Moreover, as in the case of most amino acid biosynthesis, the first step (catalyzed by anthranilate synthase) is subject to allosteric feedback inhibition by tryptophan the end product of the pathway (Kwak et al., 1999; Rouhbakhsh et al., 1996, 1997; Spraggon et al., 2001). Previous studies on different *B. aphidicola* strains have showed that all the key residues in the allosteric site are conserved in the *trpE* sequences (Lai et al., 1994; Rouhbakhsh et al., 1996). However, since the *trpE* gene of *B. aphidicola* BCc was not considered in the mentioned study, we have performed a multi-sequence analysis to compare this sequence with its homologs in other eleven *B. aphidicola* strains, and in *E. coli* and *S. marcescens.* Accordingly, we have found that the key residues P21 and S40, as well the motif LLES, purportedly involved in allosteric feedback inhibition (Caligiuri and Bauerle, 1991; Kwak et al., 1999; Tang et al., 2001), are even better conserved in *B. aphidicola* BCc than in the other *B. aphidicola* strains (see Supplementary Figure S6, and Materials and Methods sections for more details), indicating that a product inhibition mechanism could be still active.

**Figure 5.**
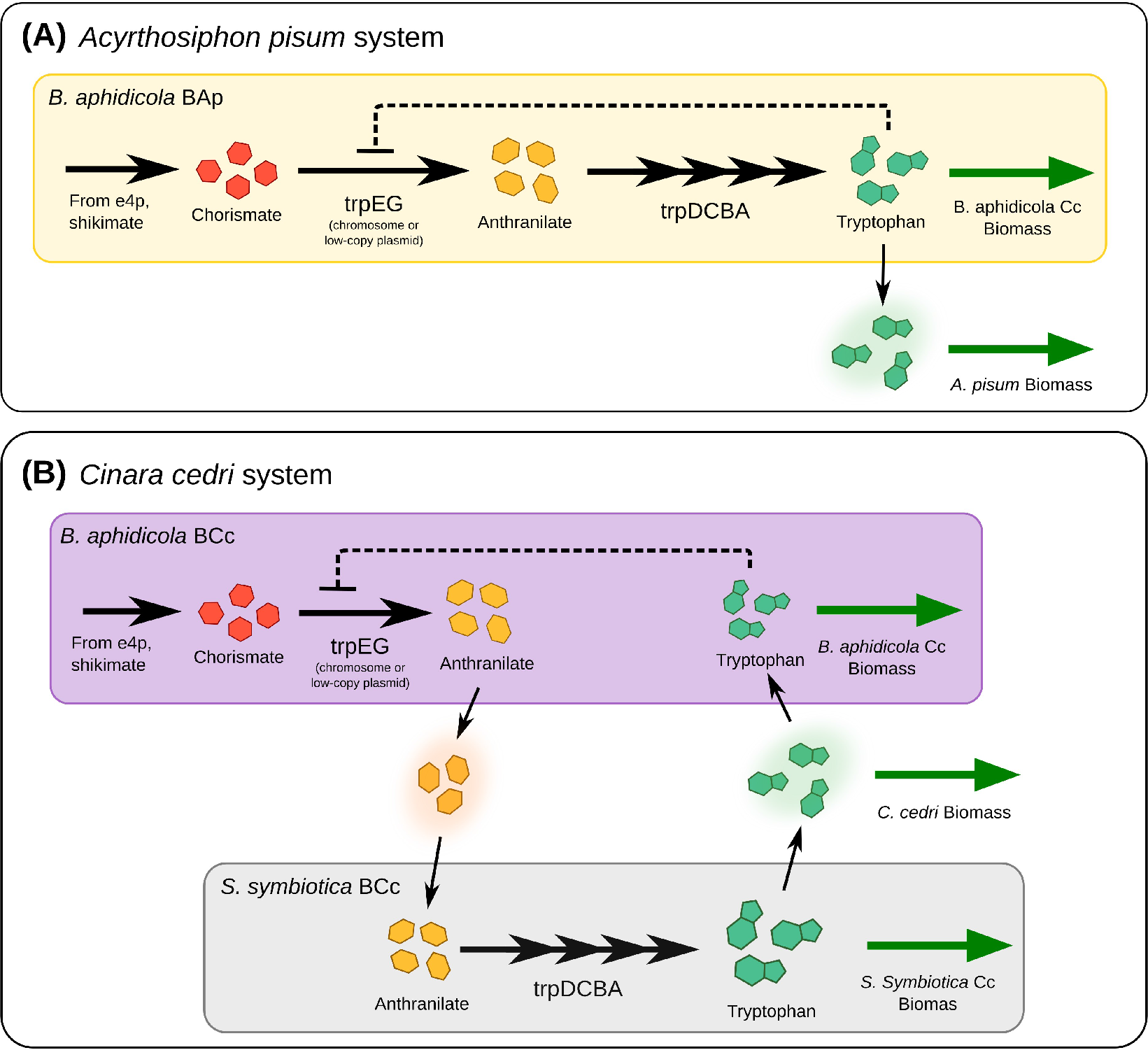
Tryptophan biosynthetic pathway in two different endosymbiont-host systems. Most Buchnera strains can synthesize tryptophan from chorismate derived from central metabolites. The first step is catalyzed by anthranilate synthase, encoded in the trpEG genes, then, anthranilate is converted into tryptophan by four additional enzymatic steps, encoded in the trpDCBA operon. In A) the structure of the pathway in the system B. aphidicola and the aphid A. pisum is presented. Feedback inhibition of tryptophan on anthranilate synthase is present. (B) In the aphid C. cedri, the primary (B. aphidicola BCc) and the coprimary (S. symbiotica SCc) endosymbionts share the production of tryptophan: trpEG genes are only present in Buchnera, while the trpDCBA genes only in Serratia.

If there is an evolutionary pressure to increase the production of tryptophan, then the inhibiting effect of tryptophan accumulation can be overcome by producing large amounts of the anthranilate synthase enzyme (Lai et al., 1994). In fact, in several *B. aphidicola* strains, *trpEG* genes are amplified, either with multiple copy of the genes in the chromosome, or with multiple-copy plasmids, suggesting an adaptation to overproduce tryptophan for hosts (Baumann et al., 2006; Rouhbakhsh et al., 1997). However, increasing protein levels to overcome the inhibition effect, may represent an large cost in terms of protein synthesis. According to our model, the metabolic complementation strategy can be used overcome feedback inhibition, while reducing the total amount of protein required to supply enough product. Indeed, in the *C. cedri* consortium the genes coding for the tryptophan biosynthetic pathway are distributed among the two different endosymbionts: the first two genes, *trpEG* encoding the anthranilate synthase activity, are hosted in a plasmid in the P-symbiont *B. aphidicola* BCc, whereas the gene products necessary to catalyze the synthesis of tryptophan from anthranilate (*trpDCBA*) are encoded by the genome of the co-primary *S. symbiotica* SCc, which in turn, lacks the *trpEG* genes (Gosalbes et al., 2008; Lamelas et al., 2011). Thus, we argue that the presence of metabolic complementation in these bacterial endosymbionts could be interpreted as an evolutionary innovation to reduce the protein cost required to supply tryptophan to all the members of the consortium.

Furthermore, the model also predicts that when a metabolic complementation is an optimal strategy, the point where the pathway should be split will correspond to the most permeable metabolite. To contrast this prediction to the present case of study, we have collected structural data of all the intermediate metabolite of the tryptophan biosynthesis pathway, and we used different physicochemical properties and rule-based estimators to evaluate membrane permeability (see the Material and Methods section), including the molecular weights and the solubility of the compounds. Using these parameters we identify plausible candidate metabolites for becoming breaking points in the tryptophan biosynthetic pathway. As seen in Table 1, among all the intermediate metabolites in this pathway, both anthranilate and indole appear as most plausible “breaking points”, while the other intermediates are less likely to cross membranes at an appreciable rate. Indole is an important signaling molecule in organisms such as *E. coli* (Martino et al., 2003), which is known to diffuse easily through cell membranes (Piñero-Fernandez et al., 2011). This raises the question: why the pathway breaking point is anthranilate instead of indole? An answer may lie in the structure of the tryptophan synthase complex (Dunn, 2012; Miles, 2013). The complex is a tetramer α2-β2. The α subunit, encoded by *trpA* gene, catalyzes the synthesis of indole from indole-3-glycerol-phosphate. Then, indole is directly channeled to the β subunit, encoded by *trpB,* where it is used to synthesize tryptophan. Interestingly, it has been shown experimentally that if the α subunit is absent, the catalytic activity of tryptophan synthase is reduced by 1-2 orders of magnitude (Kirschner et al., 1991). Therefore, keeping both *trpA* and *trpB* in the same compartment appears to be a constraint and thus, indole is not a feasible candidate for the breaking point, since it is not allowed to diffuse freely in the cell. Anthranilate is thus the intermediate most likely to diffuse through cell membranes.

**Table 1.**
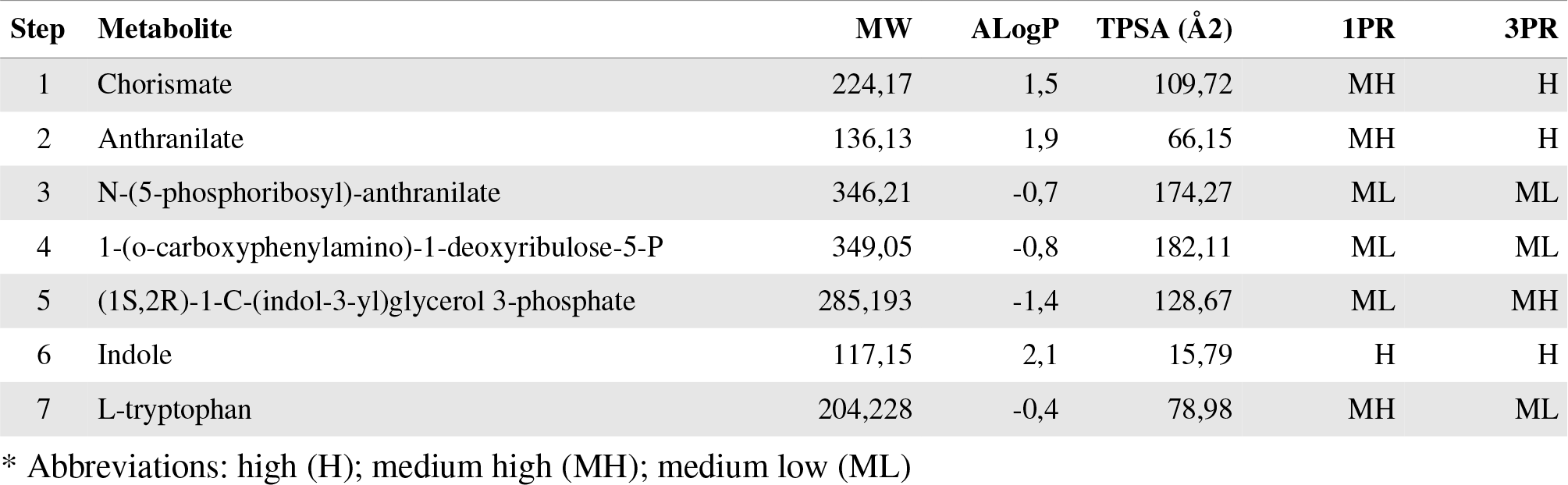
Physicochemical properties and rule-based estimators of the permeability for the intermediate metabolites of the tryptophan biosynthetic pathway. For each metabolite the molecular weight, an octanol/water logratio estimator (AlogP), the topological polar surface area (TPSA), and two rulebased classifiers (1PR and 3PR) are shown (see Estimation of metabolites permeability in Material and Methods section for details).

Finally, we will briefly discuss the case of the transaminase activities losses in different lineages of *B. aphidicola* (Hansen and Moran, 2011; Macdonald et al., 2012; Poliakov et al., 2011), as well other endosymbionts, such as *Tremblaya* and *Portiera* (Baumann et al., 2006; McCutcheon and von Dohlen, 2011; Zelezniak et al., 2015; Zientz et al., 2004). Transaminase activities are a critical step in the biosynthesis of amino acids; transamination corresponds to the last step in the production of branched-chain amino acids (BCAs) and aromatic amino acids. Moreover, all BCAs share the same transaminase enzyme (EC 2.6.1.42), and the same happens for the aromatic amino acids (*i.e.* phenylalanine and tyrosine) (with enzyme EC 2.6.1.57). In several lineages of endosymbionts, such as most strains of *Buchnera,* these transamination steps are lost, while the rest of the pathway is retained. Transaminases are expressed in the host cells (Hansen and Moran, 2011; Russell et al., 2013) originally with a catabolic function but now recruited to perform the last step in a biosynthetic pathway. As the in the case of tryptophan, the synthesis of these amino amino acids is commonly regulated at different levels, which usually include product feedback inhibition (White et al., 2012).

The above mentioned complementation does not involve two different endosymbionts, but is instead a metabolic relationship between the endosymbiont and its host. Still, this case includes all the main features of the model discussed in this work, *i.e.* a metabolite demand, feedback inhibition, and the absence of transporters. Thus, in order to test if the breaking point of these pathway corresponds to the most permeable metabolite, we have computed the permeability estimators for the intermediates in the biosynthesis of the branched amino acid as well the aromatic amino acid. The results show that in all the cases, the breaking point correspond to the most permeable metabolite (see Supplementary Table S2), thus suggesting that the complementation may have emerged due to an evolutionary pressure on the consortium.

## Discussion

The emergence of metabolic complementation in endosymbiotic consortia is a complex phenomena poorly understood from a theoretical point of view. For this reason we have developed a simple model to analyzed the possible advantages (or disadvantages) of metabolic complementation, as well as the necessary conditions that may drive its emergence. Under the model assumptions, we have found that for metabolic complementation to emerge, two necessary conditions should be fulfilled: first, the pathway must exhibit product inhibition, and the inhibition constant should be strong enough so that the over-production requires a high increase in the enzyme concentration. Second, some of the intermediate metabolites of the pathway should be permeable enough to diffuse across cell membrane so that it can reach the other cell type. Thus, an “indifferent, or “neutralist” strategy, with both cell populations allocating the same enzymes, is favored if the metabolites cannot easily permeate through the cell membranes,and/or if there is no substantial product inhibition on the first step of the pathway. On the other hand, if the intermediate metabolites can permeate easily through the membrane, and if product inhibition is present, metabolic complementation strategies allow to reduce the global enzyme burden. Moreover, when analyzing the case of tryptophan biosynthesis in the endosymbiotic consortium of *C. cedri,* we have found that the split point corresponds to the most permeable (and not channeled) metabolite. Thus, metabolite permeability is a plausible predictor of the splitting point of complementation events.

### Evolution of metabolic complementation

The evolution of metabolic complementation in bacterial communities is most likely shaped by different forces, many of them may acting simultaneously (Tan et al., 2015). On one side, endosymbionts are subject to a process of irreversible gene loss and genome shrinkage, known as the Muller's ratchet, due to its confinement to intracellular life and the bottleneck caused by its small population size (Moran, 1996). Thus, under this scenario, the gene losses with no detrimental effect on the system could be fixed at the population level by the effect of genetic drift. In this case, metabolic complementation would not necessarily confer a selective advantage, and would be a consequence of random gene losses.

Instead, a selective explanation focuses on the cost of protein synthesis, which affects both the microbial cells and the host (which also has to provide nutrients to the endosymbionts). Therefore, hosts whose endosymbionts are able to sustain a given demand of certain product while reducing the protein cost will exhibit an improvement in their fitness (Shachrai et al., 2010). The same can apply to the fitness of microbial communities as a whole; interestingly, experimental results on synthetic cross-feeding bacteria has showed that the interactions that emerge in this scenarios confer a significant fitness advantage to the genotypes involved in the interaction, and thus, stabilize the coexistence of cross-feeding organisms (Germerodt et al., 2016; Pande et al., 2014).

Another evolutionary mechanism for the emergence of metabolic complementation is given by the Black Queen Hypothesis (BQH). According to the BQH,whenever a metabolic function is redundant among the member of a microbial community, and the final product of the process leaks or escape from the producer cells, becoming a public good, the rest of the members can take advantage of this “common good” (Morris et al., 2012). Thus the BQH suggest that the loss of a leaky function will have some selective advantage at the level of each microbial species within the community; eventually, most of the members of the community will loss the genes coding for the function, until the production of public goods is just sufficient to support the community. This is not necessarily in contrast with the previous hypothesized fitness advantage for cross-feeders communities, since the loss of redundant functions may also increase the fitness of the system as a whole; the minimization of the total protein amounts might be a good approximation of the final state of a BQH-like dynamics of gene losses, where “intermediate goods” are available to all community members.

### Application to the design of synthetic communities

In perspective, understanding the evolutionary and mechanistic properties of natural microbial communities might allow the design of synthetic ones of potential industrial interest (Großkopf and Soyer, 2014; Hosoda and Yomo, 2011). For instance, the common laboratory bacteria *E. coli* have been used to study artificial communities (Hosoda et al., 2011; Pande et al., 2014). Interestingly, synthetic communities of cross-feeders based on *E. coli* mutant strains (with both over-producers and auxotrophs) were found to grow efficiently, even faster than the growth of the wild-type strain (Germerodt et al., 2016; Pande et al., 2014). Our model can be extended to these microbial communities by including the cost of protein synthesis in a genome-scale metabolic network, which has been recently been done in the context of constrained-based modeling for *E. coli* (Mori et al., 2016). However, the inclusion of allosteric regulation and diffusion still present some challenges (Machado et al., 2015; Tepper et al., 2013). This approach may eventually lead to automated design of microbial communities, which would be of extreme interest for industrial applications.

## Material & Methods

### Optimization procedure

Optimizations were solved using the *fmincon* function in Matlab 2015a for Linux. Due to the non-linear formulation of the optimization problem, the cost function exhibits multiple local minimums. Therefore, local optimization strategies such that gradient descent are not always able to find the correct minimum. However, if the parameters of the model are varied smoothly, such as in the case of the permeabilities *D_X_* and *D_P_* in Fig. 2, one may rely on the fact that the optimal solution should also vary smoothly (excluding the frontiers between the symmetric and the non-symmetric solutions). Therefore, for each combination of parameters, we optimized the cost function starting from a random seed until we got the correct minimum in all cases.

### Sensitivity analysis procedure

In the sensitivity analysis, we chose a point in the parameter space, roughly corresponding to the border between the symmetric and asymmetric region Then, we randomized by various amounts the kinetic parameters, and for each set of parameters we computed the optimal concentrations, {*x*\*}, and the corresponding optimal *E_S_* protein asymmetry, *A_S_* = ([*E*_1,*S*_] − [*E*_2,*S*_])/([*E*_1,*S*_]+[*E*_2,*S*_]), where the asterisk indicates the optimal solution. Then, for each parameters *k_i_*, we compute the following correlation: 
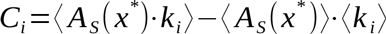
where the angled brackets ??? represent the average over the *N* samples. In order to ensure that the optimal concentration corresponded to real absolute minima of the cost function, an iterative approach was developed, which is described in detail in Supplementary Text S2. The results of the sensitivity analysis are instead in Supplementary Table S1.

### Multiple Sequence Alignment

Amino acid sequences of the anthranilate synthase component I *(trpE*) from twelve different *Buchnera* strains available, as well as *Escherichia coli* (strain K12) and *Serratia marcescens,* were downloaded from the UniProtKB database (UniProt-Consortium, 2014). The sequences in fasta format can be found in the Supplementary file Data S1. The multiple sequences alignment was performed using T-cofee package using default parameters (Notredame et al., 2000). The alignment representation was done using Jalview (Waterhouse et al., 2009).

### Estimation of metabolites permeability

A precise quantification of the diffusion constants and membrane permeability for any intermediate metabolite is not a trivial task. Therefore, we used different parameters as estimators of the permeability or diffusion capacity of the intermediate metabolites in different amino acid biosynthetic pathways we analized. For each compound we considered: (i) the molecular weight; (ii) number of hydrogen bond acceptors and donors; (iii) the octanol-water partition coefficient; and (iv) the topological polar surface area (TPSA). The structures of the amino acids biosynthetic pathways were retried from MetaCyc (Caspi et al., 2014). The list of pathways analized includes the super-pathway of aromatic amino acid biosynthesis (L-Trp, L-Tyr, L-Phe) and the super-pathway of branched chain amino acid biosynthesis (L-Leu, L-Ile, L-Val). Furthermore, all the physicochemical parameters listed above were computed using OpenBabel (O’Boyle et al., 2011), (accessed through the Python wrapper PyBel (O’Boyle et al., 2008)) with the exception of the the logarithm of the octanol-water partition coefficient (LogP) for which two different estimators were chosen, named ALogP (Tetko and Poda, 2004) and XLOP3 (Cheng et al., 2007). Results can be found in Supplementary Table S2.

## Acknowledgments

MM acknowledges financial support by Università La Sapienza of Rome, aswell as Universidad Complutense de Madrid and University of California, San Diego for technical support. MPL would like to thank Fundación Española para la Cooperatión Internacional, Salud y Polftíca Social for allowing him to carry out the research activities corresponding to this paper. Financial support from Spanish Government (grant reference: BFU2012-39816-C02-01 co-financed by FEDER funds and Ministerio de EconomÍa y Competitividad) and Generalitat Valenciana (grant reference: PR0METE0II/2014/ 065) is grateful acknowledged. MP would like to thank Martím Graña for for fruitful discussions and comments.

